# Gamma-rhythmic input causes spike output

**DOI:** 10.1101/2020.03.26.010678

**Authors:** Christopher Murphy Lewis, Jianguang Ni, Thomas Wunderle, Patrick Jendritza, Ilka Diester, Pascal Fries

## Abstract

The gamma rhythm has been implicated in neuronal communication, but causal evidence remains indirect. We measured spike output of local neuronal networks and emulated their synaptic input through optogenetics. Opsins provide currents through somato-dendritic membranes, similar to synapses, yet under experimental control with high temporal precision. We expressed Channelrhodopsin-2 in excitatory neurons of cat visual cortex and recorded neuronal responses to light with different temporal characteristics. Sine waves of different frequencies entrained neuronal responses with a reliability that peaked for input frequencies in the gamma band. Crucially, we also presented white-noise sequences, because their temporal unpredictability enables analysis of causality. Neuronal spike output was caused specifically by the input’s gamma component. This gamma-specific transfer function is likely an emergent property of in-vivo networks with feedback inhibition. The method described here could reveal the transfer function between the input to any one and the output of any other neuronal group.

## Introduction

At the heart of the brain’s processing abilities is the communication among neuronal groups. And at the heart of the brain’s cognitive functions is the flexible modulation of this neuronal communication. Numerous studies suggest that neuronal communication is affected by neuronal synchronization. For example, computational models indicate that synaptic inputs to a neuron have a higher impact when they are synchronized (Salinas and Sejnowski, 2000); intracellular neuronal recordings demonstrate reduced spike thresholds for synchronized inputs (Azouz and Gray, 2003); and recordings from awake macaque monkey reveal that neurons transmitting attended stimulus information show enhanced gamma-band synchronization (Fries et al., 2008) that in turn speeds behavioral responses (Womelsdorf et al., 2006). The communication between two neuronal groups is also affected by the synchronization between them, a concept termed Communication-through-Coherence or CTC (Fries, 2005, 2015). Computational models suggest that gamma-rhythmic inputs can entrain a postsynaptic neuronal target group and thereby enhance their impact, while at the same time reducing the impact of competing inputs (Börgers and Kopell, 2008); recordings from cats and macaques show that phase relations between neuronal groups affect their power-power covariation (Womelsdorf et al., 2007) and their transfer entropy (Besserve et al., 2015); and recordings from macaques performing selective attention tasks demonstrate that interareal gamma-band synchronization is enhanced along anatomical projections transmitting attended stimulus information (Bosman et al., 2012; Grothe et al., 2012), and that this synchronization actually improves behavioral performance (Rohenkohl et al., 2018).

Thus, recordings of neuronal activity during visual stimulation and task performance have provided strong correlative evidence for a relation between neuronal communication and neuronal gamma-band synchronization. However, whether this relation reflects a causal role of gamma has so far remained hard to test. Direct evidence for a causal role of gamma in neuronal communication might be obtained through experimental manipulation of neuronal input during concurrent measurement of neuronal spike output. We exploited the fact that synaptic input can be emulated by optogenetics. Channelrhodopsin-2 (ChR2) constitutes a light-controlled depolarizing channel in neuronal membranes, similar to transmitter-gated depolarizing channels in synapses. Yet the crucial difference is that the opening of ChR2 is under experimental control with high temporal precision. We expressed ChR2 in cat visual cortical areas 17 or 21a using adeno-associated viruses (AAVs) and Calcium/calmodulin-dependent protein kinase type II alpha chain (CamKIIα) promoters. This led to ChR2 expression almost exclusively in excitatory neurons. Thus, illumination of the infected neurons emulated synchronous excitatory inputs to groups of excitatory neurons in the investigated areas.

Stimulation with constant light confirmed previous results that cat visual cortical networks transform temporally flat input into gamma-rhythmic spike output (Ni et al., 2016). The gamma induced by constant optogenetic stimulation is very similar to gamma induced by sustained visual stimulation (Fries et al., 1997; Fries et al., 2002; Gray et al., 1989; Gray and Viana Di Prisco, 1997). We then used optogenetic stimulation to investigate how the rhythmicity of emulated input affects spike output. Specifically, we first stimulated with sine waves, as sine waves allow the modulation of frequency, while simultaneously keeping total applied light energy and peak light power constant. Sine waves of increasing frequency entrained neuronal spike responses, with a fidelity that increased up to 40 Hz and decreased only slightly for 80 Hz. Finally, and most importantly, we stimulated with white-noise sequences. In contrast to sine waves (and commonly employed regular pulse trains), white-noise is not auto-correlated, such that its causal effect on spike output can be quantified without confounds by correlated past or future input. Spike-triggered averaging of the white-noise light sequence revealed that spikes were preceded specifically by gamma-rhythmic input. Correspondingly, an analysis of Granger causality between the white-noise light input and neuronal spike output revealed a pronounced gamma-band peak.

## Results

### AAV1 and AAV9 with CamKIIα promoters transfect cat visual cortex excitatory neurons

Recombinant adeno-associated virus (AAV) vectors are widely used as gene delivery tools (Vasileva and Jessberger, 2005). AAV-mediated expression of Channelrhodopsin-2 (ChR2) has been used in several mammalian species, including mice, rats and non-human primates (Diester et al., 2011; Gerits et al., 2015; Scheyltjens et al., 2015). In this study, three pseudo-typed AAVs, AAV1, AAV5 and AAV9, were applied in visual cortex of the domestic cat (*felis catus*). We injected AAVs carrying the gene for the expression of hChR2(H134R)-eYFP under the control of the Calcium/calmodulin-dependent protein kinase type II alpha chain (CamKIIα) promoter. Injections targeted either area 17, the cat homologue of primate area V1, or area 21a, the cat homologue of primate area V4 (Payne, 1993). All AAV1 and AAV9 injections resulted in robust transfection (which was not the case for AAV5, see Methods). Transfection was evident in confocal fluorescence microscopy (and often also in epifluorescence) and in the neuronal responses to light application. In total, we transfected neurons in area 17 in four hemispheres of three cats, and in area 21a in four hemispheres of four cats.

In two cats, after electrophysiological recordings were completed, brains were histologically processed, and slices were stained for parvalbumin (PV) and/or gamma-Aminobutyric acid (GABA) (Fig. 1). One cat had been injected with AAV9-CamKIIα-ChR2-eYFP into area 21a. Across several slices and imaging windows of area 21a, we identified 182 unequivocally labeled neurons, which showed ChR2-eYFP expression or PV-anti-body staining; of those, 73 were positive for PV, and 109 expressed ChR2-eYFP, and there was zero overlap between these groups (Fig. 1A-D). The other cat had been injected with AAV1-CamKIIα-hChR2(H134R)-eYFP into area 17. Across several slices and imaging windows of area 17, we identified 284 unequivocally labeled neurons, which showed ChR2-eYFP expression or PV-anti-body staining; of those, 145 were positive for PV, and 139 expressed ChR2-eYFP, with four neurons showing clear ChR2-eYFP fluorescence and partial (patchy) PV staining (Fig. 1E-H). In the same cat, across several additional slices and imaging windows of area 17, we identified 264 unequivocally labeled neurons, which showed ChR2-eYFP expression or GABA-anti-body staining; of those, 146 were positive for GABA, and 118 expressed ChR2-eYFP, and there was zero overlap between these groups (Fig. 1I-L). Thus, ChR2 expression occurred almost exclusively in excitatory neurons.

**Figure 1.**
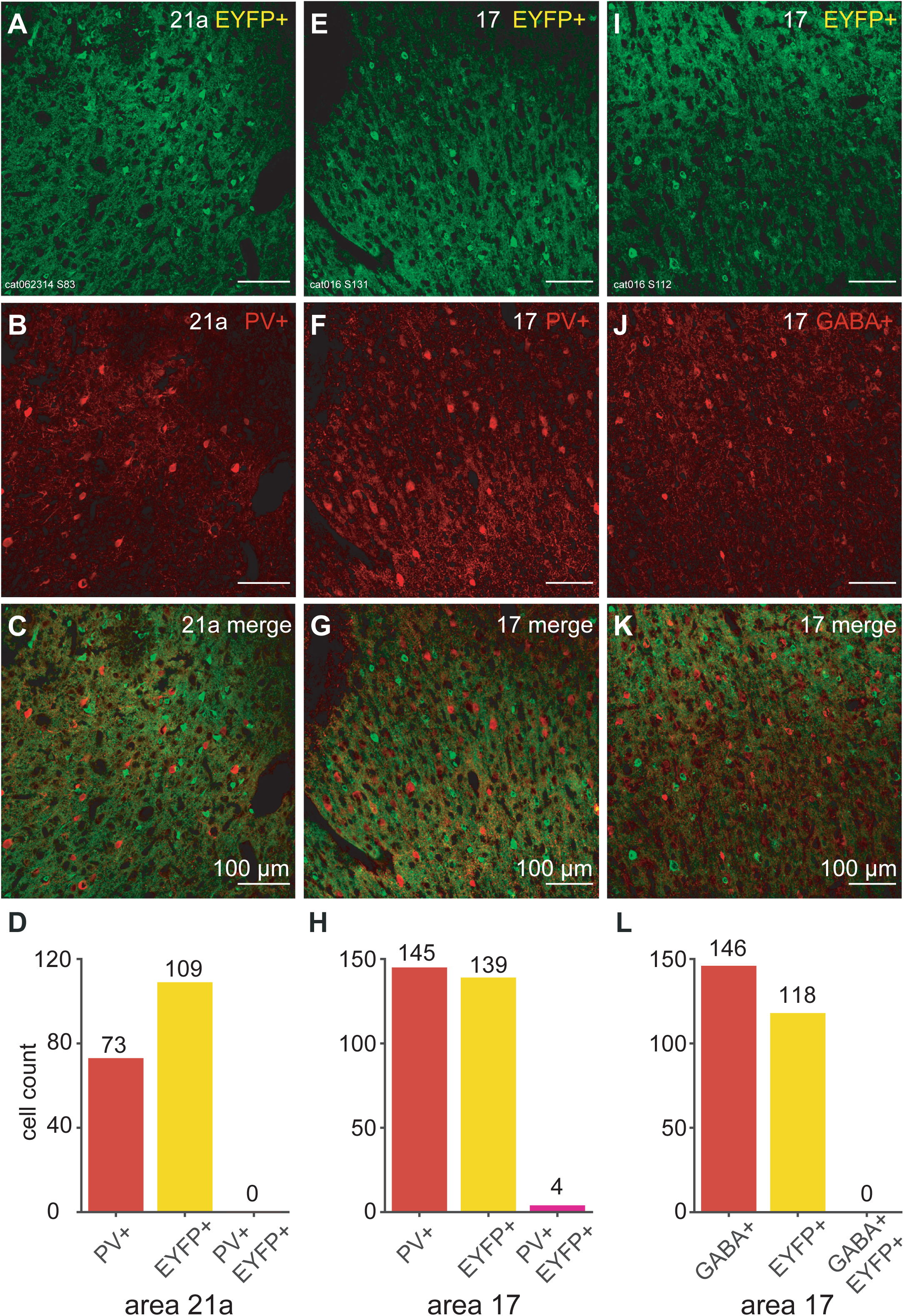
Viral transfection was largely selective for excitatory neurons. (A-C) Slice from area 21a. (E-G) Slice from area 17. (I-K) Separate slice from area 17. (A, E, I) Cells expressing ChR2-eYFP. (B, F) PV+ antibody labeled cells. (J) GABA+ antibody labeled cells. (C, G) Merged images, testing for neuronal co-labeling with ChR2-eYFP and PV+ antibody. No co-labeled neurons can be found in these slices; Four partly co-labeled neurons were found in other slices. (K) Merged images, testing for neuronal co-labeling with ChR2-eYFP and GABA+ antibody. No co-labeled neurons can be found. (D) Counts of PV+ labeled neurons, EYFP+ labeled neurons, and co-labeled neurons in area 21a. (H) Counts of PV+ labeled neurons, EYFP+ labeled neurons, and co-labeled neurons in area 17. (L) Counts of GABA+ labeled neurons, EYFP+ labeled neurons, and co-labeled neurons in area 17.

### Constant optogenetic stimulation induces neuronal gamma-band synchronization

Between 4 and 6 weeks after virus injection, we performed terminal experiments under general anesthesia. The transfected part of cortex was illuminated with blue light, while neuronal spike and LFP activity was recorded from the optogenetically stimulated region. ChR2 is a light-activated cation channel, and thereby the application of light to a neuronal group emulates excitatory synaptic inputs that are modulated according to the light modulation. Visual cortex shows particularly strong gamma-band activity in response to visual stimulation that is sustained and devoid of temporal structure (Kruse and Eckhorn, 1996). Thus, optogenetic stimulation of visual cortex might also be particularly suited to induce gamma if it is constant. Indeed, we have previously reported, in the context of an investigation of the gain-modulating effect of gamma (Ni et al., 2016), that constant optogenetic stimulation induces gamma-band activity in anesthetized cat visual cortex. Here, we present a more detailed analysis of this phenomenon. Figure 2A shows an example LFP recording from area 21a of a cat transfected with AAV9, during one trial of optogenetic stimulation with 2 s of constant blue light. The raw LFP trace reveals strong optogenetically induced gamma, and the zoomed-in epoch illustrates that this emerged immediately after stimulation onset. Figure 2B shows the spike responses of this recording site for many interleaved trials of stimulation with blue or yellow light, confirming the selective optogenetic stimulation by blue light. Figure 2C shows the spike-triggered average of the LFP, demonstrating that spikes were locked to the LFP gamma-band component. The time-frequency analysis of both, LFP power (Fig. 2D, E) and MUA-LFP locking (Fig. 2F, G) showed a strong and sustained gamma-band peak for stimulation with blue light, that was absent for stimulation with yellow light.

**Figure 2.**
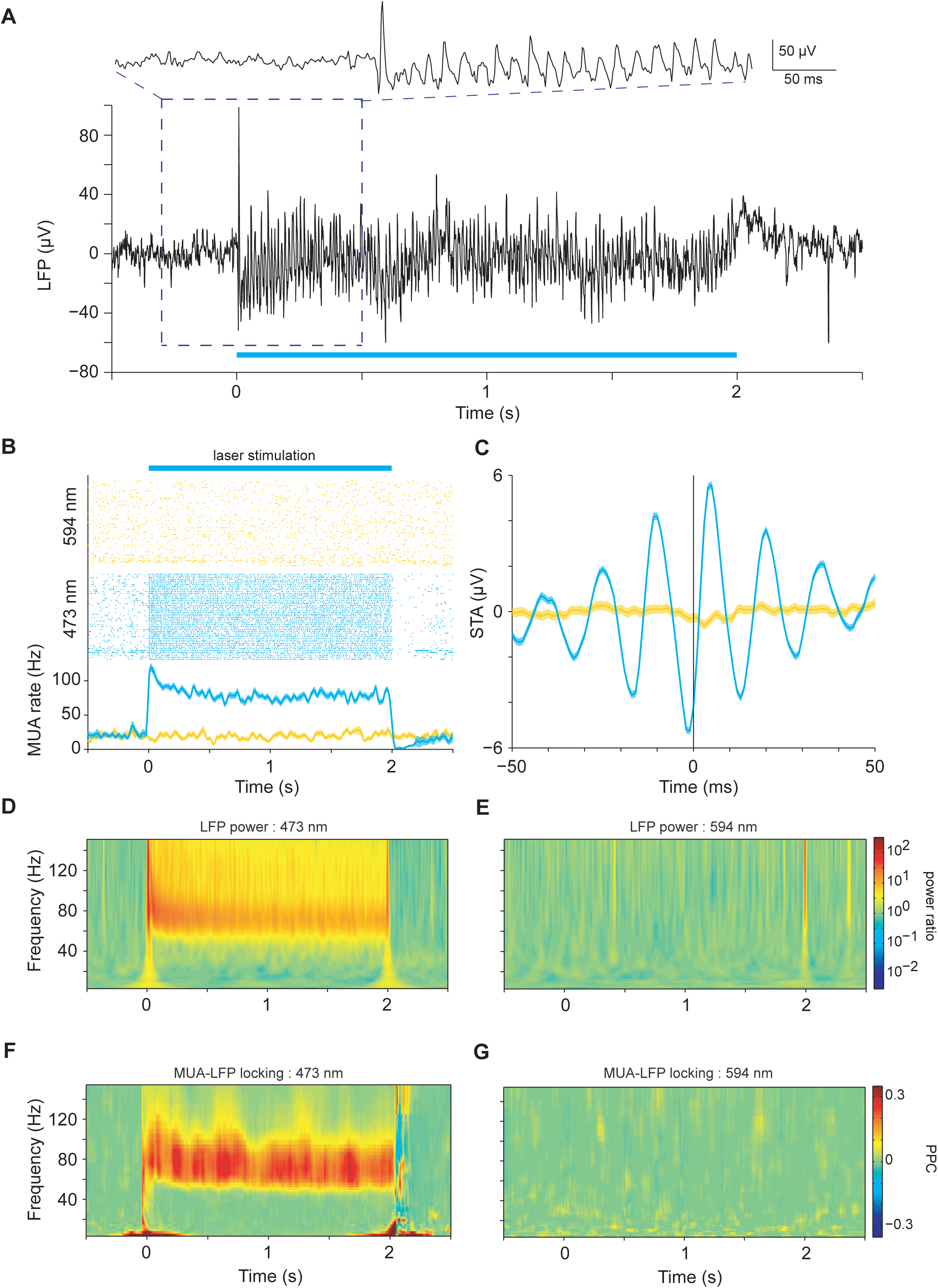
Constant optogenetic stimulation induced gamma-band synchronization. The data in this figure are from one example recording site in area 21a. (A) Constant-light induced gamma-band oscillation in the local field potential recorded in an example trial. (B) Constant-light induced MUA response. Blue: 473 nm wavelength light; Yellow: 594 nm wavelength light. The spike density was smoothed with a Gaussian function with σ = 12.5 ms, truncated at ±2*σ*. Shaded area indicates ±1 SEM. (C) Spike-triggered LFP. Shaded area indicates ±1 SEM. (D) TFR of constant-light induced gamma power, for 473 nm light. (E) Same as (D), but for 594 nm light as control. (F) TFR of constant-light induced MUA-LFP coherence. (G) The same as in (F), but for 594 nm light as control.

This pattern was found very reliably across recording sites. Stimulation with two seconds of constant blue light, as compared to yellow control light, induced strong enhancements in firing rate, which were sustained for the duration of stimulation (Fig. 3A,D; Wilcoxon rank-sum test = 39581, *p*<0.0001, n =163 sites). The ratio of LFP power during stimulation versus pre-stimulation baseline showed an optogenetically induced gamma-band peak around 70 Hz (Fig. 3B,E; Wilcoxon rank-sum test = 14751, *p*<0.0001, n = 99 sites). The LFP gamma-power changes reflected changes in neuronal synchronization, because optogenetic stimulation also induced strong MUA-LFP locking in the gamma band, as quantified by the MUA-LFP PPC (Fig. 3C,F; Wilcoxon rank-sum test = 9389, *p*<0.0001, n = 84 sites). In addition to the induction of gamma-band activity, optogenetic stimulation also caused a reduction of LFP power between 4 and 14 Hz (Fig. 3B left panel for lower frequencies; Fig.3 E inset), and a reduction in MUA-LFP locking between 10 and 12 Hz (Fig. 3C and Fig.5F inset). Those lower-frequency effects are reminiscent of effects of visual stimulation and/or selective attention in awake macaque area V4 (Fries et al., 2008).

**Figure 3.**
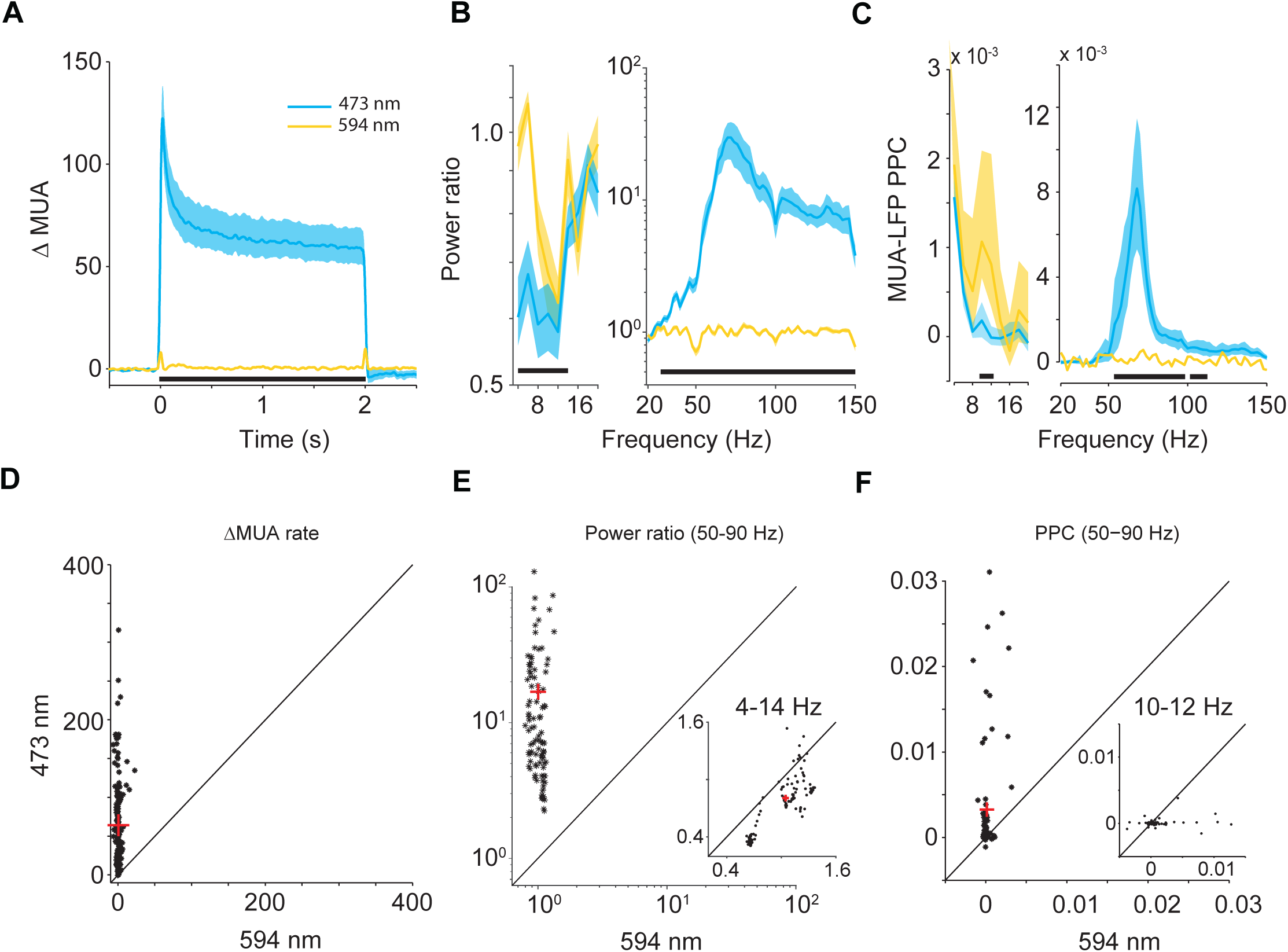
Group results for MUA rate, LFP power and MUA-LFP PPC induced by constant light. (A) Average MUA spike density. Smoothed by a Gaussian function (σ = 12.5 ms, truncated at ±2*σ*). (B) Average LFP power ratio spectrum. (C) Average MUA-LFP PPC spectrum. (B, C) use ±0.5 s epochs for the analyses from 4 to 20 Hz, and ±0.25 s long epochs for the analyses from 20-150 Hz. (A-C) Blue (yellow) lines show data obtained with 473 nm (594 nm) light stimulation. Shaded areas indicate ±1 SEM across recording sites, which is shown for illustration only. Black bars at the bottom indicate frequency ranges with statistically significant differences (*p* < 0.05), based on a cluster-level permutation test including correction for the multiple comparisons across frequencies. For the main clusters from panels (A-C), panels (D-F) illustrate the underlying distributions as scatter plots. (D) Each dot shows the MUA rate (0.3-2 s after light onset) of one recording site for blue light on the y-axis versus yellow light on the x-axis. The red cross corresponds to the respective median values. (E) Same as (D), but for the LFP power ratio (light stimulation versus pre-stimulation baseline). The main plot is for the gamma band (50-90 Hz); the inset plot for the low-frequency cluster from (B) (4-14 Hz). (F) Same as (D), but for MUA-LFP PPC during light stimulation. Each dot corresponds to one MUA recording site. The main plot is for the gamma band (50-90 Hz); the inset plot is for the low-frequency cluster from (C) (10-12 Hz).

**Figure 4.**
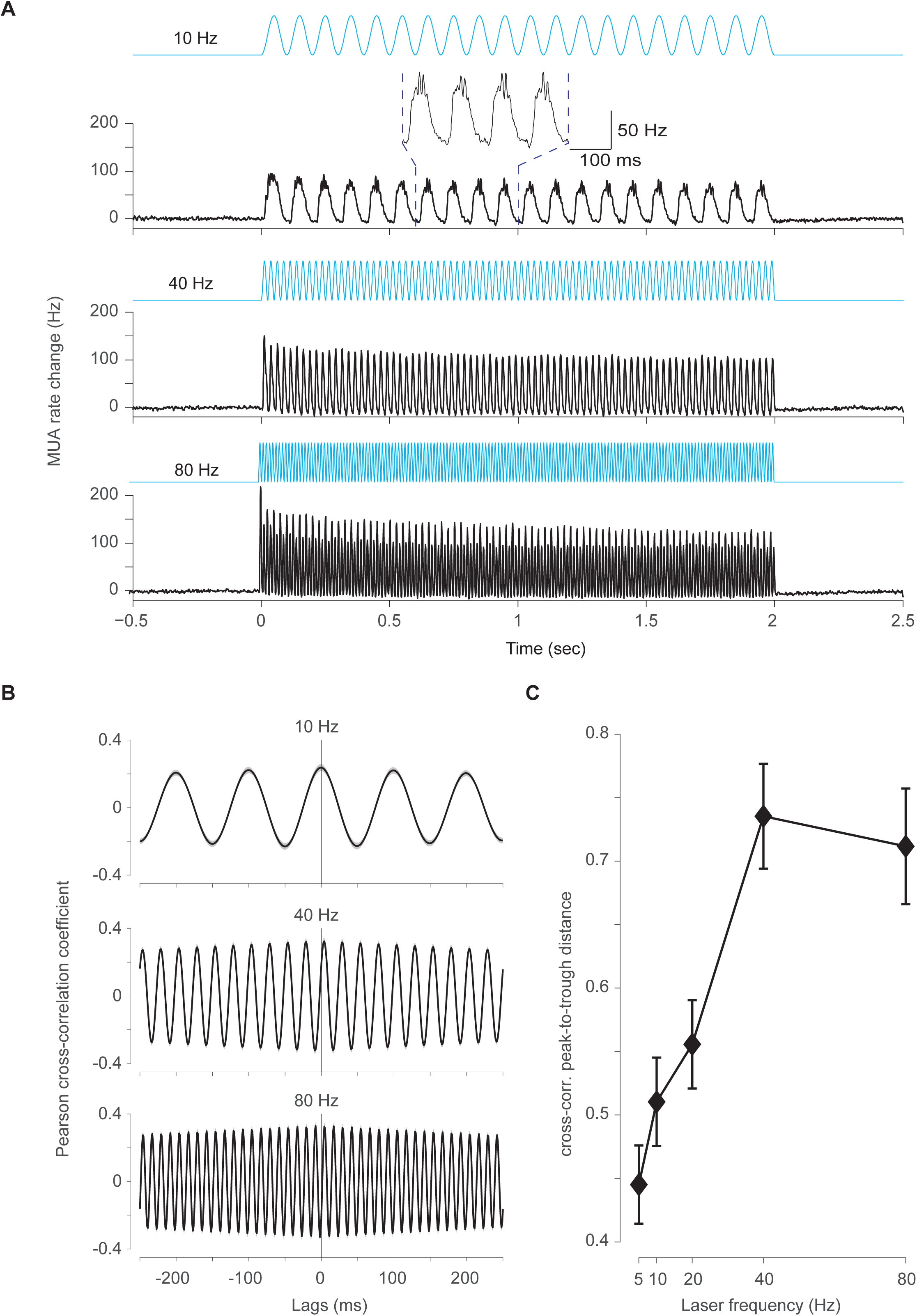
MUA responses to optogenetic sine wave stimulation. (A) MUA spike density (Gaussian smoothing with σ = 1.25 ms and truncated for ±2*σ*) for 10 Hz (top), 40 Hz (middle) and 80 Hz (bottom) optogenetic sine wave stimulation, respectively. The inset shows an enlarged version of a few cycles to illustrate the gamma-band component riding on the depolarizing phase of the 10 Hz sine wave. Data were baseline (−0.5 to 0 second) subtracted and were averaged over all MUA recording sites (N = 60). Error regions show ±1SEM across recording sites, but are barely visible. (B) Pearson cross-correlation coefficient between the analog stimulation sine wave and the MUA spike density for 10 Hz, 40 Hz and 80 Hz, respectively. (C) Modulation depth, quantified as peak-to-trough distance of the Pearson cross-correlation coefficient as function of the sine-wave frequency of optogenetic stimulation.

**Figure 5.**
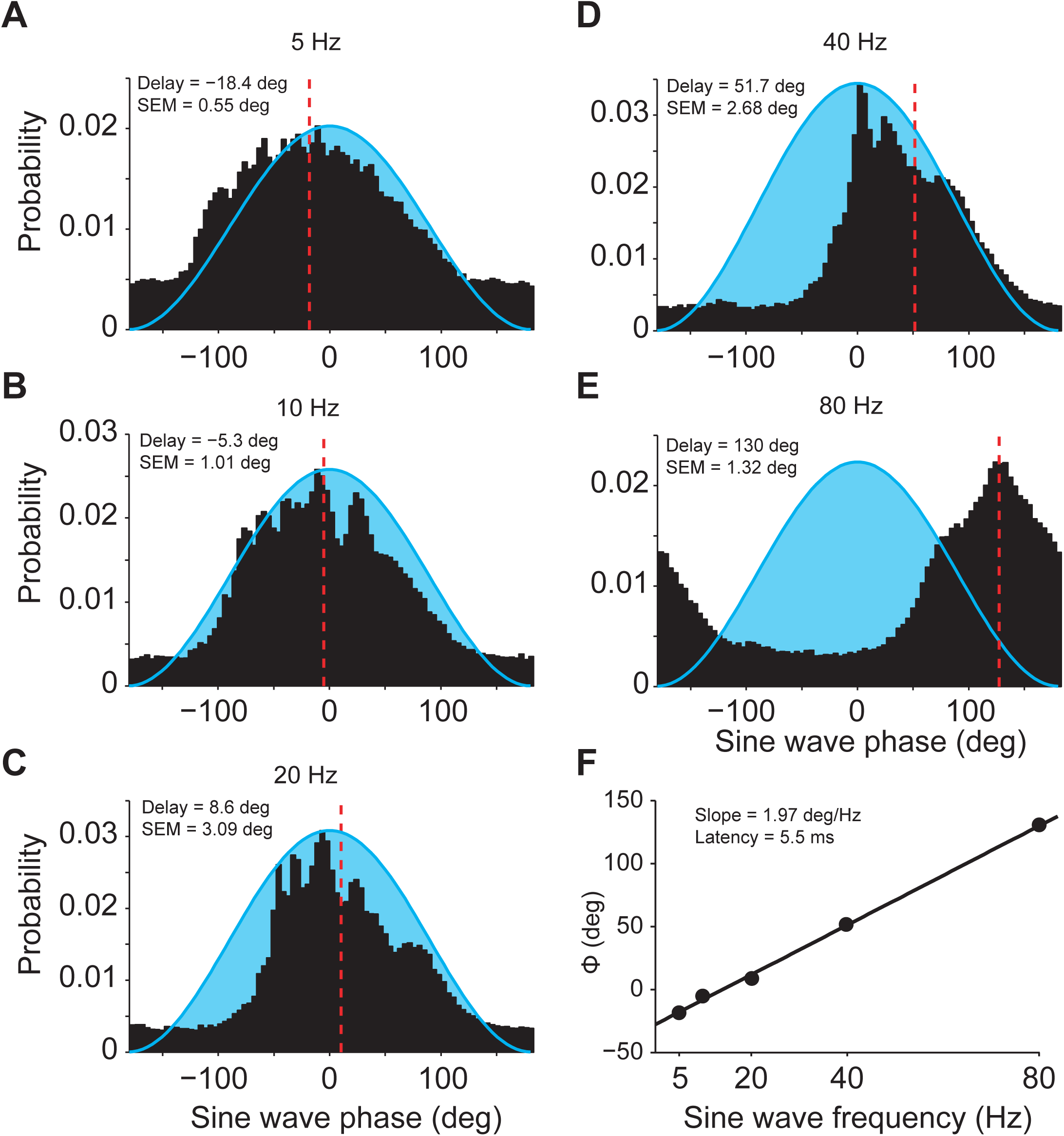
MUA response latencies to optogenetic sine-wave stimulation. (A-E) MUA spike probability, averaged over recording sites, as a function of the phase of the optogenetic sine wave stimulation. The optogenetic sine wave is indicated by the blue-shaded region. Each panel shows the data obtained with the frequency indicated on top of the panel. MUA responses were fitted with Gaussians, and the resulting peak latencies are indicated by dashed red lines. Peak latencies and their SEM (estimated through a jackknife procedure) are also indicated as text insets. Latencies are expressed relative to the time of peak light intensity. (F) MUA peak latencies from (A-E) as a function of the sine-wave frequency. The text inset gives the slope and the corresponding latency between optogenetic stimulation and neuronal response.

### Neuronal responses to optogenetic sine wave stimulation at different frequencies

We then investigated whether the spike output of a neuronal group depends on the frequency of the emulated input. We applied sine waves of 5, 10, 20, 40 and 80 Hz, similar to a previous study in cerebellum (El-Shamayleh et al., 2017). We chose sine waves for this particular investigation, rather than the more commonly used pulse trains, because they allow frequency to be varied, while simultaneously keeping total applied light energy and peak light power constant. Light intensity was adjusted per recording site (see Methods) and was kept constant for a given site across the different sine-wave frequencies. Sine waves of all employed frequencies resulted in clear increases in firing rate, with strong rhythmicity at the sine-wave frequency (Fig. 4). We calculated spike density functions, subtracted the baseline values and averaged them across recordings sites. Figure 4A shows those average spike densities for 10 Hz, 40 Hz and 80 Hz. Note that 10 Hz stimulation evoked not merely a 10 Hz following response, but also higher-frequency oscillations around each stimulation peak, in agreement with a previous report in rodent hippocampus (Butler et al., 2016). Note also that 80 Hz stimulation evoked not a simple 80 Hz following response, but that every second cycle of this 80 Hz following response was stronger, constituting a prominent 40 Hz sub-harmonic that was stable for the entire 2 s stimulation period. To capture entrainment by the optogenetic stimulation, we calculated the Pearson cross-correlation coefficient between the respective sine wave and the resulting spike density, as a function of time lag between the two (Fig. 4B). We quantified the strength of entrainment by quantifying the modulation depth of the cross-correlation functions as their peak-to-trough distance (Fig.4C). Optogenetic sine-wave stimulation resulted in entrainment that increased with stimulation frequency to peak at 40 Hz and to only weakly decline at 80 Hz (one-way ANOVA, *p* = 1.6E-9, *F*_(4,295)_ = 11.25).

Stimulation with sine waves of different frequencies allowed the estimation of neuronal response latencies. This is highly relevant when optogenetic stimulation is used to produce temporal activation patterns at high frequencies. In addition, it provides a signature of true optogenetic stimulation: Neuronal response latencies to optogenetic stimulation have typically been found to be on the order of 3-8 ms; By contrast, spikes detected at shorter latencies are suspicious of reflecting photo-electric artifacts (Cardin et al., 2010). To investigate response latencies, we averaged MUA responses aligned to the peaks of the sine waves (Fig. 5A-E). During sine-wave stimulation, the light was modulated between the respective maximal intensity and almost zero intensity. Thus, the light crossed the threshold for effective neuronal stimulation at an unknown intensity, and it is not possible to calculate response latencies in the same way as has been done for pulse trains. Therefore, we used a technique of latency estimation that has been developed in the study of synchronized oscillations, and that is based on the slope of the spectrum of the relative phase between two signals (Schoffelen et al., 2005), in our case the light intensity and the MUA. Figure 5F shows this relative-phase spectrum and reveals a strictly linear relationship of relative phase on frequency. A linear frequency-phase relation is a signature of a fixed time lag, because a given time lag translates into increasing phase lags for increasing frequencies. The slope of this linear relationship allowed us to infer a latency of 5.5 ms, which is in good agreement with previous reports of neuronal latencies.

### Optogenetic white-noise stimulation reveals causal role of gamma

Finally, and crucially, we emulated input with a white-noise characteristic. White noise realizes continuously unpredictable values (innovation), and thus shows no auto-correlation i.e. no correlation with its own past or future. Thereby, time-lagged correlations between the optogenetically emulated neuronal input and the neuronal spike output cannot be due to time-lagged correlation in the input, but can be unequivocally attributed to a time-lagged correlation between input and output. A time-lagged correlation between an experimentally controlled input and the observed spike output provides direct evidence for a causal role of the input. Importantly, white-noise stimulation enabled us to determine the causal roles separately for each frequency of the spectrum. That is, the emulation of white-noise input during recording of spike output allowed us to determine the directed transfer function of the observed network.

We employed optogenetic stimulation with light intensities following a Gaussian random process (sampled at ≈1000 Hz) with a flat power spectrum (Figure 6A, bottom trace). This white-noise stimulus contains the same energy at all frequencies up to 500 Hz. Light intensities were titrated such that firing rates were in the lower half of the dynamic range of the recorded neurons in response to optogenetic stimulation. Figure 6A shows an example LFP and MUA recording for an example trial of white-noise stimulation.

**Figure 6.**
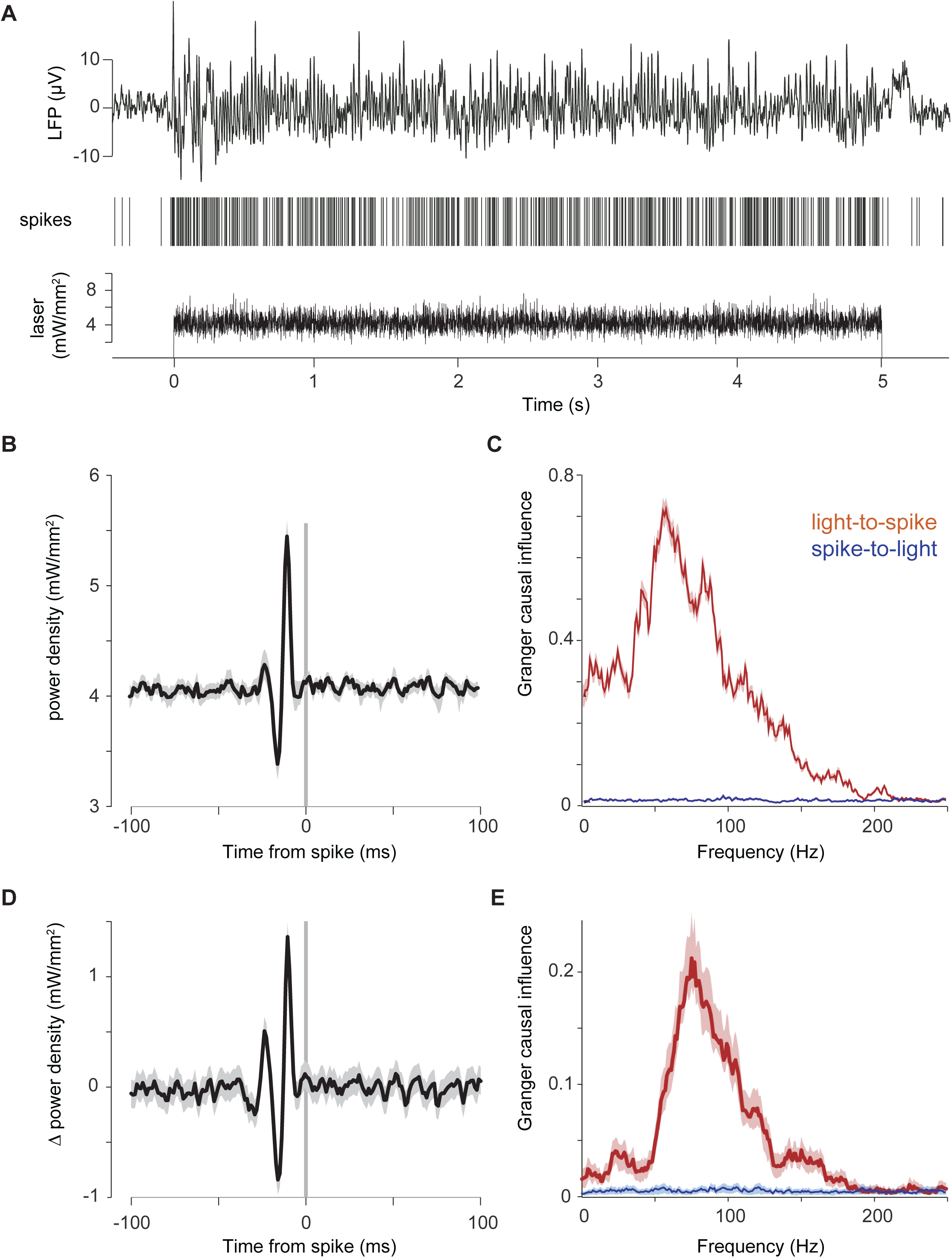
Gamma-rhythmic input causes spike output, as revealed by optogenetic white-noise stimulation. (A) Example single trial LFP and MUA response to optogenetic white-noise stimulation. The bottom panel shows the white-noise time course of laser intensity. The sequence of vertical lines above it indicates time points of MUA spike occurrence. The black continuous line on top shows the LFP. (B) Spike-triggered average (STA) of laser power density, triggered by the spikes recorded at one example recording site. (C) Granger causality (GC) spectrum for the data shown in (B). Red line shows GC from light to spikes, blue line shows GC from spikes to light (as control). (D, E) Same as (B, C), but averaged over all recording sites, for which this experiment was conducted (N=13, from 3 cats).

To reveal temporal input patterns causing spikes, we used the spikes to trigger the averaging of the light intensity. Figure 6B shows the resulting spike-triggered average light power density for an example recording site. We found that spikes were preceded by a characteristic sequence of increased and decreased light intensity, with a peak-to-peak cycle length corresponding to 75 Hz, suggesting a causal role of the gamma band in eliciting spikes. To quantify this causal influence in a frequency-resolved manner, we calculated Granger causality (GC) between the time-varying light intensity and the spike train. This revealed a clear peak in the gamma band (Fig. 6C). As a control, we also calculated the Granger causal influence of the spike train onto the light, which confirmed values close to zero, as expected. We found very similar effects in the average over recording sites (N=13) and animals (N=3) (Fig. 6D,E), confirming a predominant role of the gamma band in causing spikes.

## Discussion

We recorded LFPs and neuronal spike output in anesthetized cat visual cortex, and used optogenetics to emulate synaptic input to excitatory neurons with millisecond experimental control over the input time course. This allowed us to present input that followed a white-noise sequence, and thereby to directly quantify the input components that were causal for spike output. Spike output was specifically caused by the input’s gamma band component. This gamma-dominated input-output function agreed with the spike output that we observed in response to input that followed sinewaves of different frequencies. Sinewave inputs were translated into spike output with a fidelity that increased with frequency up to 40 Hz and only slightly declined for 80 Hz. Interestingly, when we emulated temporally flat input devoid of any structure, the stimulated visual cortical network produced spike output that was rhythmically synchronized primarily in the gamma band, as shown previously (Ni et al., 2016), and very similar to gamma induced by visual stimulation. Thus, visual cortical networks are predisposed to turn temporally flat input into gamma-rhythmic spike output, and the resulting gamma-rhythmic input to subsequent postsynaptic neurons is ideally suited to cause them to spike. This suggests that the gamma rhythm is an important aspect of visual cortical communication.

The observed input-output function for white-noise input characterizes the complete transfer chain from the signal that controlled the laser, to the spike output of the recorded neurons, which includes the laser, the opsin, the dendrites, the soma, and the action potential initiation site. Regarding the transfer function between the computer-generated control signal for the laser and the actual laser light output, we assured prior to the experiments that this was linear for the range of laser intensities used here, and that this held for the update frequency of the white noise (≈1000 Hz). Thus, the opsin molecules were stimulated by a light time course with a truly white spectrum. However, subsequent stages received input that had essentially been filtered by the transfer functions of the preceding stages. For example, the dendrites likely entailed a low-pass filtering. Such filtering could not be compensated by respective normalization of the laser control signal, because e.g. the compensation of a dendritic low-pass would have required extremely strong laser power in the dendritic stop band, which would have led to tissue damage. Further, the transfer from laser to axon hillock involves the combined effects of several distinct components, such as, multiple dendrites, and the corresponding multi-component transfer function could not be compensated by one normalization applied to the laser spectrum. Importantly, such compensation was not necessary, as we aimed to characterize the physiologically relevant compound transfer function from the emulated synaptic input to the measured spike output.

This compound transfer function showed a strong peak in the gamma band, which could be due to intracellular mechanisms, network mechanisms, or both. The intracellular transfer function has been determined in a previous study (Tchumatchenko et al., 2013) that expressed different opsin types in cultured cortical cells, applied continuously varying light with a broad spectrum of frequencies to the entire neuron, and measured currents at the soma of pyramidal cells. For the specific opsin used here (hChR2(H134R)), the observed transfer function peaks at 3 Hz, with a smooth decline by less than 10 % for lower frequencies, and a smooth decline for higher frequencies that halves currents at around 40 Hz (ChR2R in Fig. 1 of (Tchumatchenko et al., 2013)). Importantly, this transfer function does not show a peak in the gamma band. Similarly, a study that measured the transfer function between colored-noise current injected into pyramidal somata in-vitro and the resulting spike output reported resonance at 5-10 Hz and attentuation at higher frequencies (red curve in Fig. 4F of (Hasenstaub et al., 2005)). Thus, the gamma-band peak observed in the present study is most likely not due to the opsin or electrical properties of the individual neurons, i.e. it is most likely not due to intracellular mechanisms in the optogenetically stimulated excitatory neurons.

Rather, the difference between the low-frequency dominated transfer function of isolated pyramidal neurons and the gamma-band dominated transfer function of the intact in-vivo networks recorded here suggests that this gamma-band characteristic reflects a network mechanism. The network was activated by the white-noise stimulation for an extended period, similar to natural physiological conditions. Under those conditions, the recorded neurons receive excitatory and inhibitory inputs that are driven by the recent past of the light stimulation, and this includes strong perisomatic feedback inhibition with characteristic delays and decay times (Buzsáki, 2006; Buzsáki and Wang, 2012). These characteristic times for feedback inhibition likely play a central role in generating the gamma peak in the transfer function. The white-noise input sequence will occasionally contain a transient that will trigger spikes in the stimulated excitatory neurons. Those spikes will drive inhibitory neurons that will in turn cause feedback inhibition onto the excitatory neurons. When this inhibitory feedback declines with its characteristic time constant, the stimulated neurons will return to be particularly susceptible for input. Thus, inhibitory feedback renders the neuronal network particularly susceptible to input bouts spaced such that the second bout avoids the feedback inhibition triggered by the first bout, and that it overlies with the rebound excitation. Thus, feedback inhibition renders the neuronal network particularly susceptible to gamma-rhythmic input. This network mechanism is likely assisted and amplified by intracellular mechanisms in interneurons (Cardin et al., 2009; Fellous et al., 2001). Cortical interneurons recorded in-vitro show a transfer function from injected colored-noise current to spike times that peaks in the gamma range (blue curve in Fig. 4F of (Hasenstaub et al., 2005)). This gamma peak is relatively shallow, yet together with the mentioned network mechanism might well explain the pronounced gamma peak observed here in-vivo.

Importantly, the observed gamma-rhythmic pattern in the spike-triggered average of the white-noise stimulation cannot be explained by the mere fact that the stimulation induced gamma-rhythmic neuronal spiking. Rather, it required that spikes where time locked (and thereby phase locked) to the relevant temporal pattern in the white noise. If white noise had merely induced spikes that were gamma-rhythmic but not phase-locked to the gamma component of the white noise, the spike-triggered average (STA) of white noise would have been flat. Rather, the STA showed a strong gamma-rhythmic modulation. This modulation preceded the spike, because it caused the spike, and it was absent after the spike, because the spike had no causal influence on the white noise, and because white noise has no autocorrelative structure. STAs have previously been calculated for intracellularly recorded membrane potentials (Azouz and Gray, 2008; Hasenstaub et al., 2005) and LFPs (Fries et al., 1997), and this revealed that visually induced spikes were typically phase-locked particularly strongly to gamma-rhythmic components. As membrane potentials and LFPs reflect synaptic currents (Pesaran et al., 2018), these observations are consistent with a scenario in which spikes are caused specifically by the gamma components of the synaptic inputs. However, the observations are also consistent with a scenario in which visual stimulation induced gamma-rhythmic neuronal activity reflected in both spiking and LFP, without a specific causal role of gamma-rhythmic inputs. Thus, the demonstration of gamma’s causal function was enabled here by the use of optogenetic white-noise stimulation.

The experiments reported here were limited to visual cortex. Activated visual cortex shows prominent gamma-band activity (Brunet et al., 2015; Gray and Singer, 1989; Onorato et al., 2020), and the underlying mechanisms are likely related to the ones underlying the dominant role of gamma in causing spikes. If this reasoning is correct, it predicts a similar causal role of gamma in other areas, in which clear and narrowband gamma has been described (Brown et al., 1998; Csicsvari et al., 2003; Fries, 2009; Gregoriou et al., 2009; Pesaran et al., 2002). The same reasoning predicts that spikes in other areas, in which other rhythms predominate, might be caused predominantly by the corresponding rhythm in their input. Similar differences might also exist within one cortical area, between its different layers. Superficial and deep layers in macaque areas V1, V2 and V4 show very different rhythms during activation: While superficial layers show gamma, deep layers show an alpha-beta rhythm (Buffalo et al., 2011). Rhythms can also change dynamically, e.g. according to cognitive conditions, and this might change input-output functions (Gulbinaite et al., 2019). It will be a highly interesting task for future studies to perform similar experiments as described here in different areas and/or different layers, and during different cognitive conditions. Note that the approach presented here can also be used to investigate the transfer between input to one neuronal group and the spike output of another neuronal group, with the two groups possibly residing in different layers and/or areas. With recordings at site A and stimulation at sites B and C, it might even be possible to characterize not only the spectral transfer function from B to A, but also the frequency-resolved modulatory influence of C on this transfer function. Thereby, the presented approach might provide a novel framework to study neuronal communication and the mechanisms behind its cognitive modulation.

## Methods

Eight adult domestic cats (*felis catus*; four females) were used in this study. Data from the same animals were used in a previous study (Ni et al., 2016). All procedures complied with the German law for the protection of animals and were approved by the regional authority (Regierungspräsidium Darmstadt). After an initial surgery for the injection of viral vectors and a 4-6 week period for opsin expression, recordings were obtained during a terminal experiment under general anesthesia.

### Viral vector injection

For the injection surgery, anesthesia was induced by intramuscular injection of ketamine (10 mg/kg) and dexmedetomidine (0.02 mg/kg), cats were intubated, and anesthesia was maintained with N_2_O:O_2_ (60/40%), isoflurane (∼1.5%) and remifentanil (0.3 µg/kg/min). Four cats were injected in area 17 and another four cats in area 21a. Rectangular craniotomies were made over the respective areas (Area 17: AP [0, −7.5] mm; ML: [0, 5] mm; area 21a: AP [0,-8] mm, ML [9, 15] mm). The areas were identified by the pattern of sulci and gyri, and the dura mater was removed over part of the respective areas. Three to four injection sites were chosen, avoiding blood vessels, with horizontal distances between injection sites of at least 1 mm. At each site, a Hamilton syringe (34 G needle size; World Precision Instruments) was inserted with the use of a micromanipulator and under visual inspection to a cortical depth of 1 mm below the pia mater. Subsequently, 2 µl of viral vector dispersion was injected at a rate of 150 nl/min. After each injection, the needle was left in place for 10 min before withdrawal, to avoid reflux. Upon completion of injections, the dura opening was covered with silicone foil and a thin layer of silicone gel, the trepanation was filled with dental acrylic, and the scalp was sutured.

We first tried to transfect with AAV5, because this serotype had been successfully used in many studies on different species (Diester et al., 2011). In one cat, area 17 of the left hemisphere was injected with AAV5-CamKIIα-ChR2-eYFP (titer 4*10^13^ GC/ml). However, this did not result in detectable ChR2-eYFP expression. This failure of AAV5 expression is consistent with one previous study suggesting that AAV5 is not able to provide transduction in the cerebral cortex of the cat (Vite et al., 2003). Subsequently, we tried both AAV1 and AAV9 and found robust transfection with both of these serotypes. In one cat, area 17 in the left hemisphere was injected with AAV1-CamKIIα-hChR2(H134R)-eYFP (titer 8.97*10^12^ GC/ml) and area 17 in the right hemisphere with AAV9-CamKIIα-ChR2-eYFP (titer 1.06*10^13^ GC/ml). In two cats, area 17 of the left hemisphere was injected with AAV1-CamKIIα-hChR2(H134R)-eYFP (titer: 1.22*10^13^ GC/ml). In four cats, area 21a of the left hemisphere was injected with AAV9-CamKIIα-hChR2(H134R)-eYFP (titer: 1.06*10^13^ GC/ml). The DNA plasmids were provided by Dr. Karl Deisseroth (Stanford University, Stanford, CA). AAV5 viral vectors were obtained from UNC Vector Core (UNC School of Medicine, University of North Carolina, USA); AAV1 and AAV9 viral vectors were obtained from Penn Vector Core (Perelman School of Medicine, University of Pennsylvania, USA).

### Neurophysiological recordings

For the recording experiment, anesthesia was induced and initially maintained as during the injection surgery, only replacing intubation with tracheotomy and remifentanyl with sufentanil. After surgery, during recordings, isoflurane concentration was lowered to 0.6%-1.0%, eye lid closure reflex was tested to verify narcosis, and vecuronium (0.25mg/kg/h i.v.) was added for paralysis during recordings. Throughout surgery and recordings, Ringer solution plus 10% glucose was given (20 ml/h during surgery; 7 ml/h during recordings), and vital parameters were monitored (ECG, body temperature, expiratory gases).

Each recording experiment consisted of multiple sessions. For each session, we inserted either single or multiple tungsten microelectrodes (∼1 MΩ at 1 kHz; FHC), or three to four 32-contact probes (100 µm inter-contact spacing, ∼1 MΩ at 1 kHz; NeuroNexus or ATLAS Neuroengineering). In one cat, one 16-contact probe with 150 µm inter-contact spacing and one 46 µm optic fiber, and one 16-contact probe with 150 µm inter-contact spacing and four 46 µm optic fibers were used (Plexon V- and U-probe, respectively). Standard electrophysiological techniques (Tucker Davis Technologies, TDT) were used to obtain multi-unit activity (MUA) and LFP recordings. For MUA recordings, the signals were filtered with a passband of 700 to 7000 Hz, and a threshold was set to retain the spike times of small clusters of units. For LFP recordings, the signals were filtered with a passband of 0.7 to 250 Hz and digitized at 1017.1 Hz.

### Photo-stimulation

Optogenetic stimulation was done with a 473 nm (blue) laser or with a 470 nm (blue) LED (Omicron Laserage). A 594 nm (yellow) laser was used as control. Laser light was delivered to cortex through a 100 µm or a 200 µm diameter multimode fiber (Thorlabs), LED light through a 2 mm diameter polymer optical fiber (Omicron Laserage). Fiber endings were placed just above the cortical surface, immediately next to the recording sites with a slight angle relative to the electrodes. Laser waveform generation used custom circuits in TDT, and timing control used Psychtoolbox-3, a toolbox in MATLAB (MathWorks) (Brainard, 1997).

For white-noise stimulation, the laser was driven by normally distributed white noise, with light intensities updated at a frequency of 1017.1 Hz. For each recording session, the mean of the normal distribution was chosen to fall into the lower half of the dynamic range of the laser-response curve of the recorded MUA. This resulted in mean values in the range of 3-12 mW/mm^2^ (13 MUA recording sites in area 17 of 3 cats). The standard deviation (SD) of the normal distribution was scaled to be 1/2 the mean. The resulting distributions were truncated at 3.5 SDs. The resulting range of laser intensities always excluded both zero and maximal available laser intensities and thereby avoided clipping.

### Histology

After conclusion of recordings, approximately five days after the start of the terminal experiment and still under narcosis, the animal was euthanized with pentobarbital sodium and transcardially perfused with phosphate buffered saline (PBS) followed by 4% paraformaldehyde. The brain was removed, post-fixed in 4% paraformaldehyde and subsequently soaked in 10%, 20% and 30% sucrose-PBS solution, respectively, until the tissue sank. The cortex was sectioned in 50 µm thick slices, which were mounted on glass slides, coverslipped with an antifade mounting medium, and subsequently investigated with a confocal laser scanning microscope (CLSM, Nikon C2 90i, Nikon Instruments) for eYFP-labelled neurons.

#### Immunohistochemistry

In two cats, one with injections in area 17 and one with injections in area 21a, slices were processed as described above and additionally stained for parvalbumin (PV) and gamma-Aminobutyric acid (GABA). To this end, slices were preincubated in 10% normal goat serum (NGS) with 1% bovine serum albumin (BSA) and 0.5% Triton X-100 in phosphate buffer (PB) for 1 h at room temperature to block unspecific binding sites. Floating slices were stained for PV (over night, rabbit anti-Parvalbumin, NB 120-11427, Novus Biologicals) and GABA (48 hours, rabbit anti-GABA, ABN131, Merck Millipore) in 3% NGS containing 1% BSA and 0.5% Triton X-100. After washing two times 15 min in PB, the slices were incubated with the secondary antibody (goat anti-rabbit Alexa Fluor 647, A-21244, Thermo Fisher Scientific) in 3% NGS containing 1% BSA and 0.5% Triton X-100 for 1 h at room temperature. Finally, slices were again washed in PB, coverslipped and imaged with a Zeiss CLSM, using a 25X water immersion objective.

### Data analysis

All data analysis was performed using custom code and the Fieldtrip toolbox (Oostenveld et al., 2011), both written in MATLAB (MathWorks).

#### Spike densities, MUA-laser cross-correlation, LFP power spectra, and MUA-LFP PPCs

MUA rate was smoothed with a Gaussian (for constant light stimulation: SD = 12.5 ms; for stimulation with sine waves: SD = 1.25 ms; in each case truncated at ±2 SD) to obtain the spike density.

To quantify the locking of neuronal responses to optogenetic stimulation, we calculated the Pearson correlation coefficient between MUA spike density and laser intensity as a function of time shift between them.

LFP power spectra were calculated for data epochs that were adjusted for each frequency to have a length of 4 cycles and moved over the data in a sliding-window fashion in 1 ms steps. Each epoch was multiplied with a Hann taper, Fourier transformed, squared and divided by the window length to obtain power density per frequency. For the different stimulation frequencies f, LFP power is shown as ratio of power during stimulation versus pre-stimulation baseline (−0.5 s to −0.2 s relative to stimulation onset).

MUA-LFP locking was quantified by calculating the MUA-LFP PPC (pairwise phase consistency), a metric that is not biased by trial number, spike count or spike rate (Vinck et al., 2010). Spike and LFP recordings were always taken from different electrodes. For each spike, the surrounding LFP was Hann tapered and Fourier transformed. Per spike and frequency, this gave the MUA-LFP phase, which should be similar across spikes, if they are locked to the LFP. This phase similarity is quantified by the PPC as the average phase difference across all possible pairs of spikes. To analyze PPC as a function of frequency and time (Fig. 2F), the LFP around each spike in a window of ±2 cycles per frequency was Hann tapered and Fourier transformed. PPC was then calculated for epochs of 100 ms length, i.e. using the phases of spikes in those epochs, moved over the data in a sliding-window fashion in 1 ms steps. To analyze PPC with higher spectral resolution (Fig. 3), the LFP around each spike in a window of ±0.5 s (Fig. 3C, lower frequencies; Fig. 3F, inset) or ±0.25 s (Fig. 3C, higher frequencies; Fig. 3F main plot) was Hann tapered and Fourier transformed to obtain the spike phase. For a given MUA channel, MUA-LFP PPC was calculated relative to all LFPs from different electrodes and then averaged.

#### Estimation of Granger causality (GC) between light time course and MUA spike trains

The GC spectrum was first estimated separately for each recording site and subsequently averaged over sites. For each trial, we estimated the Fourier transforms of the input (laser) and the output (MUA). Specifically, each trial was segmented into non-overlapping epochs of 500 ms length. Per epoch, the time series of the input and the output were multiplied with a Hann taper, they were zero-padded to a length of 1000 ms, and their Fourier transforms (FTs) were obtained. The FTs were used to calculate the power-spectral densities (PSDs) of the input and of the output, and the cross-spectral density (CSD) between input and output. CSDs and PSDs were averaged over trials and used for the estimation of GC by means of non-parametric spectral matrix factorization (Dhamala et al., 2008). For the example GC spectrum (Fig. 6C), the error region was determined by a bootstrap procedure, with 100 iterations, each time randomly choosing 30% of the trials. The shown error boundary is the region containing 95% of the bootstrapped estimates. For the average GC spectrum (Fig. 6E), the error region indicates the standard error of the mean across the recording sites.

#### Statistical testing

All inferences were based on the combined data of all animals, for which a given experiment was performed. The resulting inferences are limited to the studied sample of animals, as in most neurophysiological in-vivo studies.

High-resolution spectra of LFP power changes and MUA-LFP PPC were compared between stimulation with blue light and control stimulation with yellow light (Fig. 3B,C). We calculated paired t-tests between spectra obtained with blue and yellow light, across recording sites. Statistical inference was not based directly on the t-tests (and therefore corresponding assumptions will not limit our inference), but the resulting t-values were merely used as a well-normalized difference metric for the subsequent cluster-based non-parametric permutation test. For each of 10,000 permutations, we did the following: 1) We made a random decision per recording site to either exchange the spectrum obtained with blue light and the spectrum obtained with yellow light or not; 2) We performed the t-test; 3) Clusters of adjacent frequencies with significant t-values (p<0.05) were detected, and t-values were summed over all frequencies in the cluster to form the cluster-level test statistic. 4) The maximum and the minimum cluster-level statistic were placed into maximum and minimum randomization distributions, respectively. For the observed data, clusters were derived as for the randomized data. Observed clusters were considered significant if they fell below the 2.5^th^ percentile of the minimum randomization distribution or above the 97.5^th^ percentile of the maximum randomization distribution (Maris and Oostenveld, 2007). This corresponds to a two-sided test with correction for the multiple comparisons performed across frequencies (Nichols and Holmes, 2002).

## Acknowledgements

This work was supported by DFG (SPP 1665 FR2557/1-1, FOR 1847 FR2557/2-1, FR2557/5-1-CORNET, FR2557/6-1-NeuroTMR, FR2557/7-1-DualStreams to P.F.; EXC 1086, DI 1908/5-1, DI 1908/6-1 to I.D.), BMBF (01GQ1301 to I.D.), EU (HEALTH-F2-2008-200728-BrainSynch, FP7-604102-HBP, FP7-600730-Magnetrodes to P.F.; ERC Starting Grant OptoMotorPath to I.D.), a European Young Investigator Award to P.F., the FENS-Kavli Network of Excellence to I.D., National Institutes of Health (1U54MH091657-WU-Minn-Consortium-HCP to P.F.), the LOEWE program (NeFF to P.F. and I.D.).

## Author contributions

C.M.L., J.N, T.W., P.F. designed research; C.M.L., J.N, T.W., P.J., I.D., P.F. performed experiments; C.M.L., J.N., T.W., P.F. analyzed data; C.M.L., J.N., P.F. wrote the paper.

